# Baclofen decreases compulsive alcohol drinking in rats characterised by reduced levels of GAT-3 in the central amygdala

**DOI:** 10.1101/2020.06.29.178236

**Authors:** Lucia Marti-Prats, Aude Belin-Rauscent, Maxime Fouyssac, Mickaёl Puaud, Paul J. Cocker, Barry J. Everitt, David Belin

**Affiliations:** Department of Psychology, University of Cambridge, UK

**Author notes:** Correspondence to: Dr. David Belin, (Dpt. of Psychology, University of Cambridge, Downing street, Cambridge, CB2 3EB, UK).

**Keywords:** Alcohol, Baclofen, Central amygdala, Compulsive alcohol drinking, GABA transporter GAT-3

## Abstract

While most individuals with access to alcohol drink it recreationally, about 5 % lose control over their intake and progressively develop an alcohol use disorder (AUD), characterised by compulsive alcohol drinking accompanied by decreased interest in alternative sources of reinforcement. The neural and molecular mechanisms underlying the vulnerability to switch from controlled to compulsive alcohol intake have not been fully characterized, so limiting the development of new treatments for AUD. It has recently been shown that rats having reduced levels of expression of the gamma-aminobutyric acid (GABA) transporter, GAT-3, in the amygdala tend to persist in seeking and drinking alcohol even when adulterated with quinine, suggesting that pharmacological interventions aimed at restoring GABA homeostasis in these individuals may provide a targeted treatment to limit compulsive alcohol drinking. Here, we tested the hypothesis that the GABA_B_ receptor agonist baclofen, which decreases GABA release, specifically decreases compulsive alcohol drinking in vulnerable individuals. In a large cohort of Sprague-Dawley rats allowed to drink alcohol under an intermittent two-bottle choice procedure, a cluster of individuals was identified that persisted in drinking alcohol despite adulteration or the availability of an alternative ingestive reinforcer, saccharin. In these rats, that were characterised by decreased GAT-3 mRNA levels in the central amygdala, acute baclofen administration (1.5 mg/kg, intraperitoneal) resulted in a decrease in compulsive drinking. These results indicate that low GAT-3 mRNA levels in the central amygdala represent an endophenotype of AUD and that the associated compulsive alcohol drinking characteristic is sensitive to baclofen.

## 1 Introduction

About 2.3 billion people worldwide consume alcohol^1^. While most maintain control over their recreational drinking, a subset of individuals are vulnerable to progressively develop alcohol use disorder (AUD)^2^, characterised by compulsive alcohol seeking and drinking that persists despite adverse consequences, and to the detriment of other activities and sources of reinforcement^3-5^. Given the major health burden that AUD places on our society, estimated to cause 7.2% of the premature deaths worldwide^1^, a better understanding of the neural, cellular and molecular basis of the vulnerability to shift from recreational to compulsive alcohol drinking may enable novel treatment targets to be identified.

Many preclinical and clinical studies have highlighted the important involvement of the gamma-aminobutyric acid (GABA) system in the regulation of the reinforcing properties of alcohol and in the development of AUD. Pharmacological manipulations of GABAergic transmission through targeting both GABA_A_ and GABA_B_ receptors, influence different behavioural responses to alcohol, including alcohol self-administration, voluntary alcohol drinking, relapse-like and binge-like alcohol drinking, and the motivation for alcohol^6-10^. Furthermore, alterations in the expression of genes encoding some GABA_A_ and GABA_B_ receptor subunits, as well as GABA transporters, have been shown to influence the predisposition to high alcohol consumption and vulnerability to AUD in humans, as well as AUD-like behaviour in rodents^11, 12^.

The central nucleus of the amygdala (CeA) is a key neural locus in which the GABAergic system mediates neuroadaptations involved in alcohol drinking behaviours^13, 14^. Both acute and chronic alcohol exposure increase GABAergic transmission in CeA and alcohol-dependent rats show increased baseline GABA levels in the CeA compared to alcohol-naïve rats^13, 14^. Infusion of a GABA_A_ receptor antagonist into the CeA reduces alcohol self-administration in non-dependent rats^15^, whereas microinjection of a GABA_A_ receptor agonist reduces self-administration in alcohol-dependent but not in non-dependent rats^14, 16^, indicating that the development of alcohol dependence is associated with adaptations in the GABAergic system within the CeA.

Beyond its involvement in controlling alcohol drinking and alcohol dependence, the GABAergic system within the CeA has also been shown to be involved in the development of addiction-like behaviours for alcohol. Augier et al^17^ recently revealed a down-regulation of the mRNA level for several genes involved in GABA homeostasis in a subpopulation of rats that both preferred alcohol over an alternative reinforcer in an exclusive-choice instrumental procedure, and also showed the addiction-like behaviours^18, 19^ of a high motivation for alcohol and the persistence of drinking when alcohol was adulterated with the bitter tastant, quinine^17^. Down-regulation of the GABA transporter GAT-3 in the amygdala was also shown to be associated both with the choice of alcohol over an alternative ingestive reward and with compulsive alcohol drinking. GAT-3 down-regulation was also found in the CeA of postmortem brains of individuals having been diagnosed with AUD^17^. Together, these data strongly suggest that alterations in the expression of GAT-3 could be a relevant candidate in the search of markers of vulnerability to, and also a treatment target for AUD.

Functionally, GABA transporters mediate GABA uptake and control extracellular GABA levels and GABAergic transmission in the central nervous system^20, 21^, and inhibition of the GABA transporters results in an increase of extracellular levels of GABA^20^. This is consistent with the finding that rats having a propensity to develop compulsive alcohol seeking and drinking that are characterised by decreased expression of GAT-3, also show increased GABAergic tone in CeA^17^. This indicates that restoring the putatively altered extracellular GABA levels triggered by reduced GAT-3 expression may decrease persistent alcohol drinking in compulsive individuals^17, 22^. For example, GABA receptor agonists such as baclofen may restore extrasynaptic GABA levels by acting at presynaptic GABA_B_ receptors that control GABA release in the synaptic cleft^22-24^ and hence decrease alcohol drinking.

Activation of GABA_B_ receptors has been shown to prevent preparatory and consummatory responses for alcohol, such as, the acquisition and maintenance of voluntary alcohol drinking, binge-like alcohol drinking and instrumental responding for alcohol^6,8-10^. Adaptations to chronic alcohol exposure such as the alcohol deprivation effect and the reinstatement of the extinguished alcohol seeking behaviour are also prevented by GABA_B_ receptor activation^6, 8^. However, the influence of GABA_B_ receptor agonists on compulsive alcohol drinking behaviour, a characteristic of AUD, has not yet been investigated.

Here, we hypothesised that GABA_B_ agonists may rescue GAT-3-related deficits in GABA homeostasis associated with the vulnerability to develop compulsive alcohol drinking. We therefore tested the effects of systemically administered baclofen on alcohol drinking in rats identified as compulsive or resilient to AUD following a long history of exposure to alcohol under an intermittent two-bottle choice procedure.

The results show that rats characterised by lower GAT-3 mRNA levels in the CeA tended to persist in drinking alcohol despite the availability of an alternative ingestive sweet reward and adulteration with quinine, two behavioural features of AUD^3,25-27^. We further show that while baclofen dose-dependently decreased voluntary alcohol intake in all rats, it selectively decreased the quinine-resistant, compulsive, drinking of alcohol in vulnerable rats.

## 2 Materials and Methods

### Subjects

Forty-eight adult male Sprague-Dawley rats (Charles River Laboratories, Kent, UK) weighing approximately 250 g on arrival were single-housed in a temperature- (22 ± 1 °C) and humidity- (60 ± 5%) controlled environment, under a 12-h reverse light/dark cycle (light on at 7:00 PM) with ad libitum access to food (standard chow) and water. After two weeks of habituation to the vivarium and prior to be given access to alcohol, rats were progressively food restricted to reach 85–90% of their free-feeding body weight. All procedures were conducted under the project licence number PPL 70/8072, in accordance with the United Kingdom Animals (Scientific Procedures) Act 1986, amendment regulations 2012 following ethical review by the University of Cambridge Animal Welfare and Ethical Review Body (AWERB).

### Experimental procedures

After two weeks of habituation to the animal facility followed by two weeks of food restriction, rats (n=48) were given intermittent access to alcohol in a two-bottle choice procedure^28, 29^. One individual failed to acquire alcohol drinking and was therefore excluded from all analyses. After three weeks of intermittent access to alcohol in the two-bottle choice procedure^28, 29^, the compulsive alcohol drinking was assessed over successive challenge sessions during which the persistence of drinking alcohol was measured when alcohol was adulterated with quinine (session 10)^25,26,30,31^ or in the presence of an alternative reward^17,27,32^, saccharin (session 14). These two tests were separated by three baseline sessions. Previous studies have shown that exposure to saccharin prior to, or after a prolonged history of alcohol exposure does not influence the tendency to choose one over the other^17^.

Adulteration-Alternative reward Resistant, Intermediate and Sensitive phenotypes (AAR, AAI and AAS, respectively) were characterised based on the persistence in drinking alcohol in both tests by a k-mean cluster analysis^33-35^. Subsequently, rats were re-baselined in the intermittent two-bottle choice procedure (alcohol vs water) over three weeks and habituated to receive intraperitoneal (IP) administrations with saline. Then, the effect of baclofen on voluntary alcohol intake was measured over three sessions following a Latin square design with one baseline session between each administration (sessions 23 to 27). The population received IP injections of saline (0), baclofen 1 mg/kg (1) and 1.5 mg/kg (1.5) (n=47/dose) 30 minutes prior to the beginning of the two-bottle choice session^10^. Thereafter, rats were re-baselined for nine weeks prior to be subjected to a single alcohol adulteration test (session 54) during which the effect of baclofen on compulsive drinking was tested in a between-subject design. Each AAR and AAS subpopulation was split into two groups each, matched for their level of alcohol intake at baseline, which received IP administration of saline (AAR, n=13; AAS, n=8) or baclofen 1.5 mg/kg (AAR, n=10; AAS, n=9) 30 minutes before starting the two-bottle choice session on which alcohol was adulterated with the bitter tastant quinine. Rats were re-baselined for three weeks (up to session 62) prior to being sacrificed and their brains harvested for the subsequent molecular analysis. GAT-3 mRNA levels were measured both in CeA and basolateral amygdala (BLA) of rats from both AAS and AAR groups. One CeA and two BLA samples were not included in the final analysis due to technical issues, so that the final sample size was AAR: n=22 and n=21 vs AAS: n=17 and n=17 for CeA and BLA, respectively (**Figure 1**).

**Figure 1:**
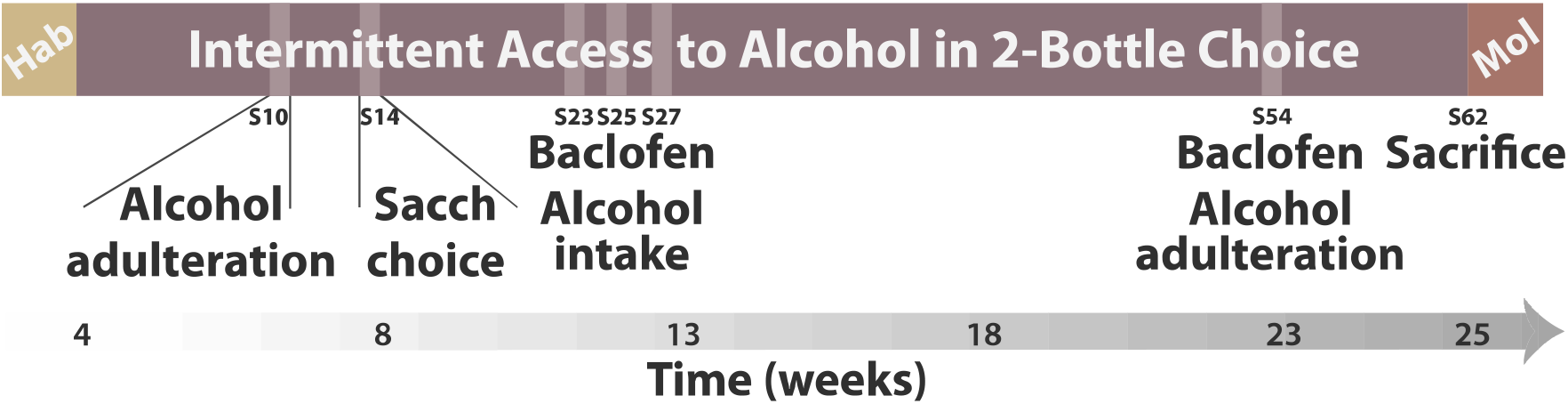
Timeline of the experiments. After four weeks of habituation to the vivarium during which they were progressively food deprived, male Sprague Dawley rats were given intermittent access to a 10% alcohol in two-bottle choice procedure. After three weeks of alcohol exposure, the compulsive nature of drinking behaviour was assessed over successive challenge sessions during which the propensity of each individual to persist in drinking alcohol despite adulteration with quinine (session 10) or an alternative reward, e.g. saccharin (session 14) was measured. Three subpopulations, namely Adulteration-Alternative reward Resistant (AAR), Intermediate and Sensitive (AAS), were characterised by K-means clustering. Rats were re-baselined in the intermittent two-bottle choice procedure for three weeks and the effect of baclofen (1 and 1.5 mg/kg) on voluntary alcohol drinking was measured for the entire population following a Latin square within-subjects design with a baseline session between treatments (Baclofen Alcohol intake, sessions 23 to 27). Rats then were re-baselined for nine weeks and the effect of baclofen on the persistence in alcohol drinking despite adulteration for the AAR and AAS subpopulations was assessed following a between subject design (Baclofen Alcohol adulteration, session 54). Lastly, rats were re-baselined before being sacrificed (session 62) and their brains harvested for posterior molecular analysis. Hab: habituation; Mol: molecular analysis; s: session; Sacch choice: saccharin choice.

### Drugs

10% (v/v) alcohol solution was prepared by mixing 99.8% ethanol (Sigma-Aldrich) in tap water. Quinine hydrochloride dihydrate (Sigma-Aldrich) was dissolved in 10% alcohol at the concentration of 0.1 g/L^36^. Sodium saccharin (Sigma-Aldrich) was dissolved in tap water at the concentration of 0.2% (v/v)^37^.

(±)-Baclofen (Sigma, UK) was freshly prepared, formulated at 1 and 1.5 mg/kg, dissolved in sterile 0.9% saline. Baclofen was administered IP in a volume of 1 ml/kg. The doses of baclofen were selected based on previous studies ensuring unspecific effects were avoided^9^.

### Intermittent access to two-bottle choice for alcohol

Rats were given intermittent access to alcohol under a two-bottle choice procedure^28, 29^. During each session, rats had free access to two bottles containing either 10% alcohol or water for twenty-four hours in their home-cage. During the intermitting twenty-four- or fortyeight-hour alcohol-free periods, rats had access to two bottles of water resulting in an opportunity to drink alcohol three times a week. The location of the alcohol bottle was switched between sessions to avoid any side preference bias.

The bottles were weighed before and immediately after each two-bottle choice session and the volume consumed per session was computed as the weight difference between the start and the end of the sessions minus the spillage volume, accounted for the volume of fluid lost from bottles placed on an empty cage^33, 34^.

During the alcohol adulteration tests, alcohol solution was substituted with alcohol containing 0.1g/L quinine for twenty-four hours (alcohol + quinine vs water).

During the saccharin choice test, saccharin 0.2% (v/v) was added to the water bottle for twenty-four hours (alcohol vs water + saccharin).

### Tissue collection and RNA extraction

Rats were deeply anaesthetised by 5% isoflurane inhalation and decapitated. Brains were harvested, frozen by immersion in −35/40°C isopentane for ~3 min and stored at −80°C. Bilateral samples from the CeA and BLA were obtained using a micro-puncher (1 mm) from 300 μm-thick coronal sections obtained using a cryostat and stored at −80°C. RNAs were extracted using the Quick-RNA™ Microprep kit (Zymo Research) following the manufacturer guidelines and quantified using the NanoDrop^®^ ND-1000 spectrophotometer (Thermo Fisher Scientific).

### Quantitative polymerase chain reaction (qPCR)

RNA was reverse-transcribed into cDNA with the RT^2^ First Strand Kit (Qiagen, UK) according to the manufacturer instructions. The following primers were used to assess the relative level of GAT-3 mRNA: Slc6a11 Rn.10545 in comparison to that of cyclophilin A (Peptidylprolyl isomerase A) used as housekeeping gene (Qiagen, UK). Real-time-PCR was performed on the CFX96 Real-Time PCR Detection System (Bio-Rad, UK) using the RT^2^ SYBR Green Mastermix (Qiagen, UK). The relative mRNA level of the target gene was calculated using CFX Manager Software (Bio-Rad, UK) and expressed as 2^-ΔCT^ relative to cyclophilin A.

### Data and statistical analysis

Data are presented as boxplots [median ± 25% (percentiles) and Min/Max as whiskers]. The individual propensity to persist in drinking alcohol (as % of baseline) was calculated as the ratio between alcohol + quinine (Alcohol adulteration test) or alcohol (Saccharin choice test) intake (g/kg) over the average alcohol intake during the last three pre-test baseline sessions. Statistical analyses were performed with STATISTICA 10 software (StatSoft, Inc., Tulsa, OK, USA). Assumptions for parametric analysis, namely normality of distribution, homogeneity of variance and sphericity were verified prior to each analysis with Shapiro-Wilk, Kolmogorov-Smirnov, Cochran and Mauchly’s tests, respectively.

K-means cluster analysis, carried-out as previously described^33-35^, on the persistence in alcohol drinking despite adulteration or access to saccharin, identified three specific subpopulations: AAR, AAI or AAS.

Persistence in alcohol drinking (as % of baseline) in the adulteration and saccharin choice tests was analysed with repeated-measures analysis of variance (ANOVA) with test as within-subject factors and group (AAR vs AAS) as between-subject factor. The effect of baclofen on alcohol and water intake (g/kg) on the whole population was subjected to repeated-measures ANOVA with treatment (three doses) as within-subject factors. The differential effect of baclofen on AAR and AAS rats on baseline alcohol and water intake (g/kg) or compulsive alcohol drinking (% of baseline) was analysed using a similar ANOVA with group (AAR vs AAS) or group (AAR vs AAS) and treatment (2 doses) as a between-subject factor, respectively. Total alcohol intake (g/kg), body weight and GAT-3 mRNA levels were analysed with a one-way ANOVA with group (AAR vs AAS) as between-subject factor.

Dimensional analyses were performed with a Pearson correlation analysis with alcohol intake (g/kg) and adulterated alcohol intake (g/kg) as variables on each of the AAR and AAS subpopulations.

For all analyses, upon confirmation of main effects, differences among individual means were further analysed using Newman-Keuls post-hoc test. Significance was set at α ≤ 0.05. Effect sizes are reported using partial eta-squared value (pη^2^)^35^.

## 3 Results

Following nine sessions of alcohol drinking under a two-bottle choice procedure, the persistence of alcohol drinking when adulterated with quinine, or when saccharin-sweetened water was available as an alternative reward was tested in each individual (**Figure 1**). Marked individual differences were observed in the persistence of alcohol drinking in each of the two tests that led to the characterisation by cluster analysis of three subpopulations, namely AAR, n=23; AAI, n=7 and AAS, n=17.

AAR and AAS were considered as vulnerable and resilient to AUD, respectively (**Figure 2A**), since the former were resistant to both challenges which produced a significant decrease in alcohol intake in the latter [main effect of group: F_1,38_=95.322, p<0.001, pη^2^=0.71; test x group interaction: F_1,38_=1.346, p=0.253] (**Figure 2B**). These behavioural manifestations were not attributable to a differential prior exposure to alcohol because AAR and AAS rats did not differ in their total alcohol intake before the tests [main effect of group: F_1,38_=2.6893, p=0.109] (**Figure 2C**). Nor did AAR and AAS differ in their body weights [main effect of group: F_1,38_=2.0772, p=0.158, data not shown].

**Figure 2:**
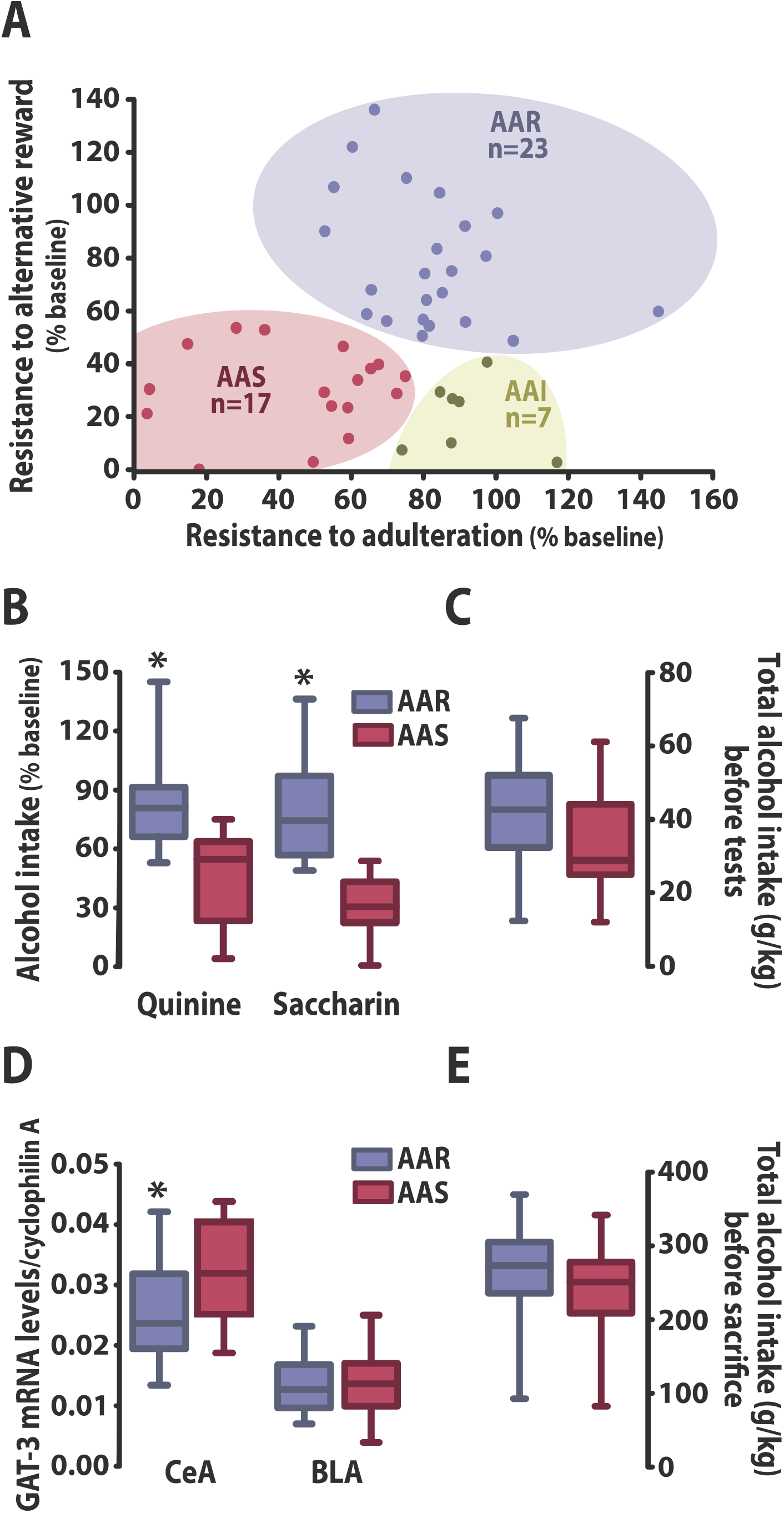
The vulnerability compulsively to drink alcohol is underlined by a low GAT-3 mRNA level in the Central Amygdala. **(A)** A cluster analysis on both the persistence in alcohol drinking despite adulteration with quinine and access to a saccharin solution, identified three subpopulations in a large cohort of outbred rats, resistant (AAR, n=23), intermediate (AAI, n=7) or sensitive (AAS, n=17) rats. **(B)** While AAS rats substantially decreased their alcohol intake when alcohol was adulterated or saccharin was offered as an alternative, the AAR subpopulation maintained high levels of alcohol intake in both tests. These differential behavioural manifestations were not attributable to a differential alcohol exposure before the tests **(C). (D)** AAR rats showed lower GAT-3 mRNA levels than the AAS rats in the central amygdala (CeA) (n=22 and n=17, respectively) (left panel) but not in the basolateral amygdala (BLA) (n=21 and n=17, respectively) (right panel). The lower GAT-3 mRNA levels shown by AAR rats in the CeA was not attributable to a differential alcohol exposure prior to sacrifice. * p<0.05 different from AAS rats. AAR: Adulteration-Alternative reward Resistant rats; AAI: Adulteration-Alternative reward Intermediate rats; AAS: Adulteration-Alternative reward Sensitive rats.

As predicted, AAR rats showed a lower level of GAT-3 mRNA in the CeA than the AAS rats [main effect of group: F_1,37_=4.564, p=0.039, pη^2^=0.11], but not in the BLA [main effect of group: F_1,36_<1] (**Figure 2D**). The decreased GAT-3 mRNA levels in the CeA shown by AAR rats was specifically related to their compulsive alcohol drinking and not due to a differential level of alcohol exposure prior to sacrifice [main effect of group: F_1,38_=2.139, p=0.152] (**Figure 2E**).

Baclofen induced, at the population level, a dose-dependent decrease in the voluntary intake of alcohol [main effect of treatment: F_2,92_=4.691, p=0.011, pη^2^=0.09] (**Figure 3A**) but not water [main effect of treatment: F_2,92_=1.515, p=0.225] (**Figure 3B**).

**Figure 3:**
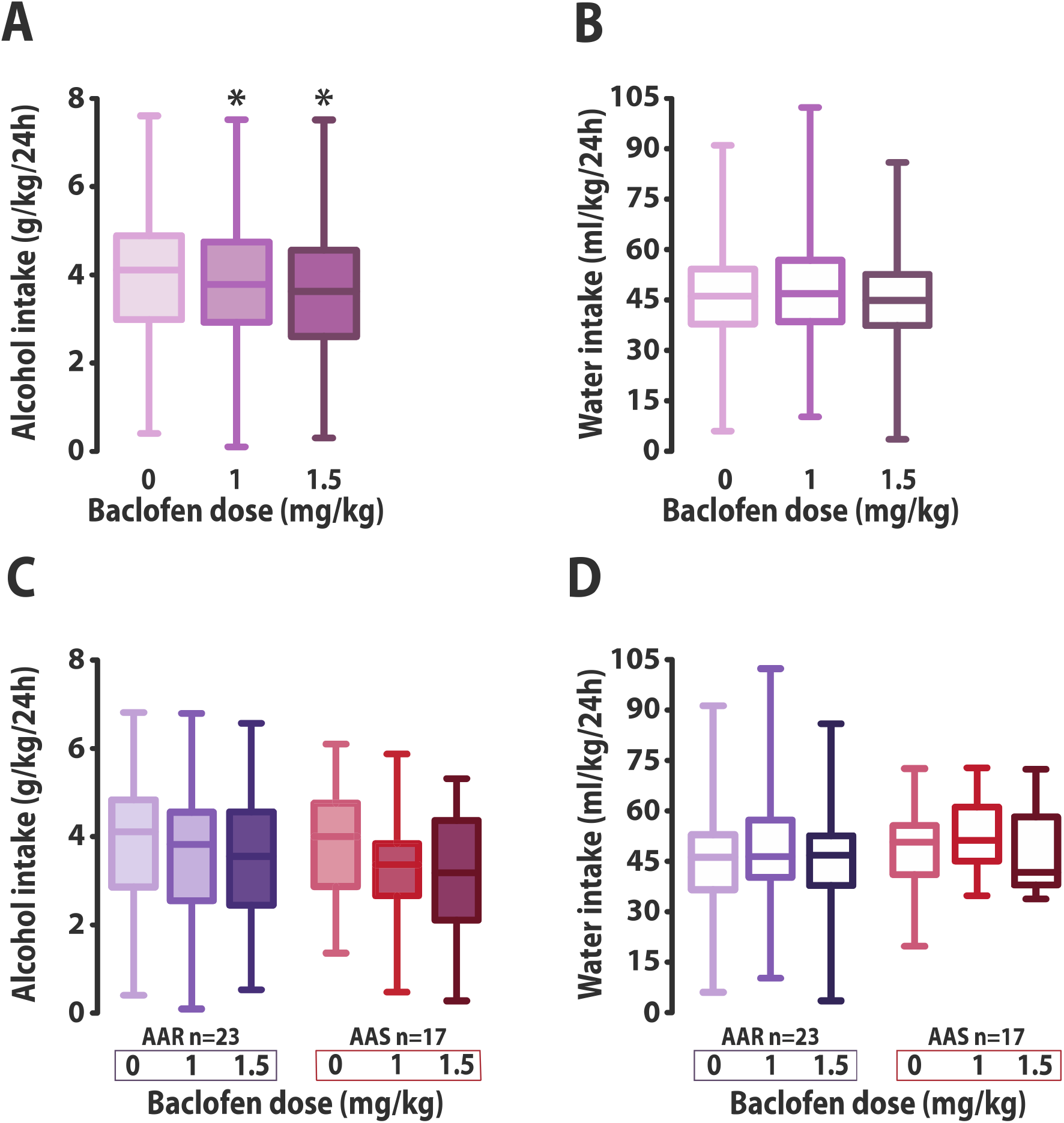
Baclofen reduces voluntary alcohol drinking in heterogeneous rat population level. Systemic administration of baclofen (1 and 1.5 mg/kg, intraperitoneal) dose-dependently reduced voluntary alcohol **(A)**, but not water intake **(B)** in the two-bottle choice procedure in a population of 47 rats. The decrease in voluntary alcohol intake after baclofen administration was not driven by either the AAR (n=23) or the AAS rats (n=17), since both groups displayed similar reduction in alcohol **(C)** while maintaining constant water intake **(D)** when treated with baclofen. * p<0.05 compared with saline treatment (0 mg/kg group). AAR: Adulteration-Alternative reward Resistant rats; AAS: Adulteration-Alternative reward Sensitive rats.

The specific reduction in baseline alcohol intake by systemic administration of baclofen was of similar magnitude between AAR and AAS rats [main effect of treatment: F_2,76_=7.655, p<0.001, pη^2^=0.17; treatment x group interaction: F_2,76_<1] (**Figure 3C**), having no effect on water intake [main effect of treatment: F_2,76_=2.370, p=0.100; treatment x group interaction: F_2,76_<1] (**Figure 3D**). In marked contrast, baclofen selectively reduced compulsive, quinine-resistant alcohol drinking shown by AAR rats when both groups where challenged with another quinine adulteration test four months later [main effect of group: F_1,36_=7.07, p=0.012, pη^2^=0.16; group x treatment interaction: F_1,36_=5.441, p=0.025, pη^2^=0.13] (**Figure 4A**). The selective sensitivity displayed by AAR rats to baclofen when they expressed persistent, compulsive drinking was not correlated with that they showed for their unchallenged drinking at baseline (R=0.5617, p=0.091), no correlation was observed for AAS rats either (R=0.2583, p=0.502) (**Figure 4B**).

**Figure 4:**
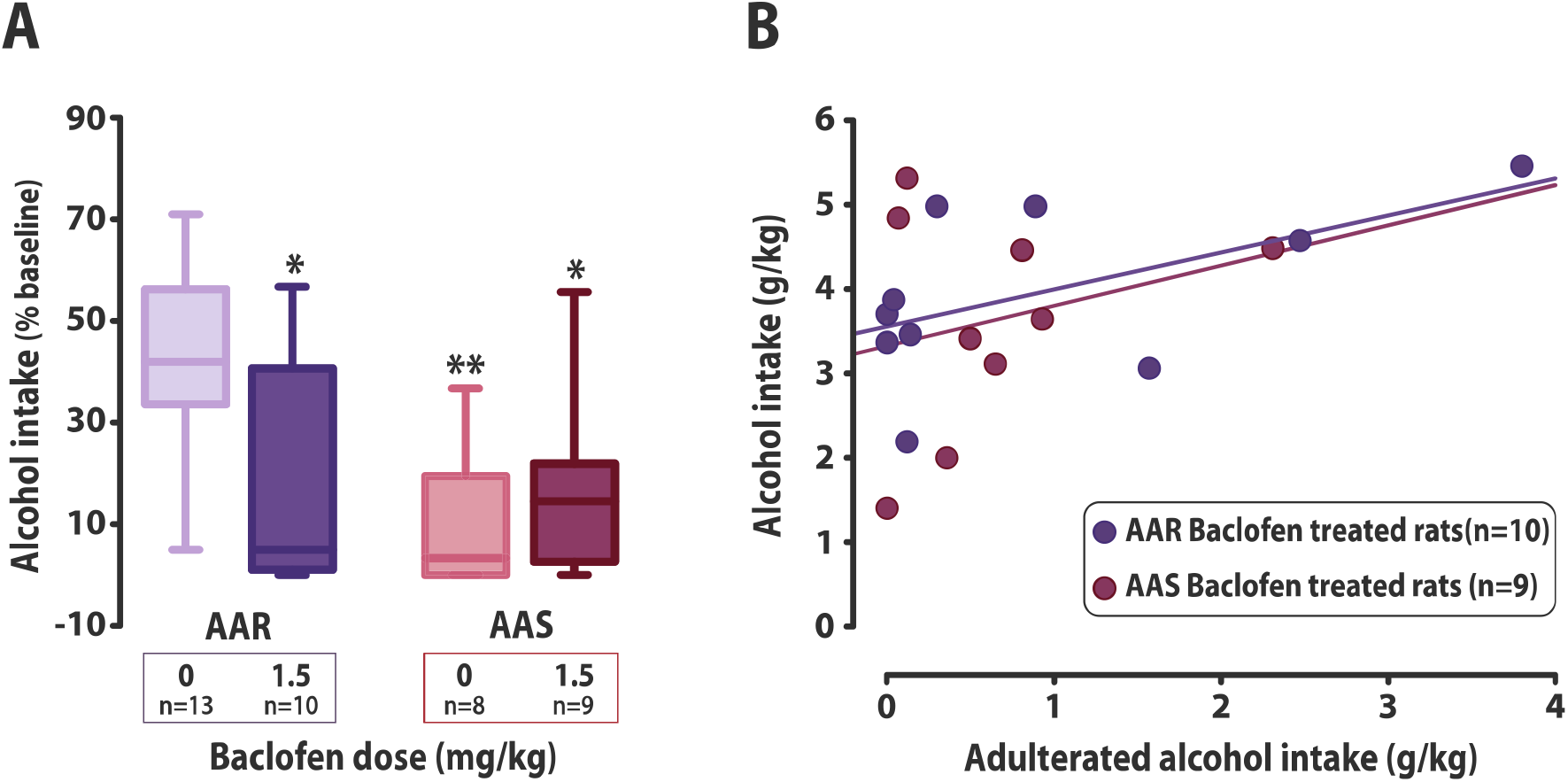
Baclofen selectively decreases persistent, compulsive alcohol drinking. **(A)** The compulsive phenotype of AAR rats was maintained after four additional months of intermittent exposure to alcohol as revealed by their persistence of drinking despite adulteration with quinine. Baclofen (1.5 mg/kg, intraperitoneal) selectively decreased compulsive drinking in AAR rats while having no influence on the alcohol drinking shown by the AAS subpopulation to be substantially decreased by adulteration. **(B)** The selective effect baclofen was shown to exert on persistent drinking in AAR rats was absolutely independent of, and unrelated to, that baclofen had on drinking under baseline conditions. *: p<0.05, **: p<0.01 compared with the saline treated (0 mg/kg) AAR group. AAR: Adulteration-Alternative reward Resistant rats; AAS: Adulteration-Alternative reward Sensitive rats.

## 4 Discussion

The results of this study reveal that following intermittent two-bottle choice exposure to alcohol, a procedure known to result in the progressive escalation of alcohol intake^28, 29^, only a subset of vulnerable individuals that were characterised by low GAT-3 mRNA levels in the CeA, went on to persistently drink alcohol when adulteration with quinine^25,26,30,38^ or when an alternative, highly preferred sweet reward was available^17,27,39^.

These resistant rats displayed behaviours that have been suggested to be the operationalisation in rodents of two features that characterise AUD, namely compulsive drinking behaviour and decreased interest for reinforcers other than the drug itself^25-27^.

In line with previous studies^26,34,40^, the development of compulsive drinking behaviour in vulnerable animals was shown not to be predicted by the individual tendency to drink high levels of alcohol, thereby confirming that the development of this behavioural feature of alcohol addiction is not simply related to the magnitude of pharmacological exposure to alcohol. This dissociation, which challenges a drug-centred view of drug addiction and further emphasising the importance of factors of vulnerability^41, 42^, is not restricted to alcohol. The propensity to switch from controlled to cocaine addiction-like behaviours in rats is also independent of the level of cocaine intoxication^18, 43^ while a history of escalated cocaine intake does not facilitate the development of compulsive cocaine self-administration^44^ or compulsive cocaine seeking^43^.

These inter-individual differences in the propensity to drink alcohol despite its adulteration or the availability of a sweet ingestive reward that emerged after three weeks of intermittent access to alcohol under a two-bottle choice procedure, persisted for at least four months. This support the construct validity of the experimental design used here and suggests that adaptations at the psychological and neural systems level that result in the emergence of compulsive alcohol drinking (present study) and seeking^33, 34^, persist over time regardless of further intoxication.

Previous studies^30-32,36^ have suggested that longer exposure to alcohol (3 to 6 months^36^) is necessary for the emergence of a quinine-resistant phenotype in rats, whereas this characteristic appeared more rapidly in the present study. However, even though the dose of quinine used here is similar to that used in the previous studies, several important methodological differences preclude any direct comparison. In the present study, quinine resistance was measured in a home-cage based non-operant procedure, as in the seminal studies of Wolffgramm and colleagues^30, 31^. However, in their intermittent conditions, alcohol was available only once a week as opposed to three times a week in the present study. On the other hand, when rats had been trained to drink alcohol under similar intermittent access conditions to those used here, the effect of quinine was not assessed on drinking under free access conditions, but on instrumental performance under a progressive ratio of alcohol reinforcement^36^. Additionally, in previous studies, the resistance to quinine adulteration was measured at the population level and did not factor-in the individual vulnerability that clearly emerged in the design of the present study.

This individual vulnerability to develop compulsive alcohol drinking behaviour was shown in here to be associated with adaptations in the GABAergic system in the CeA. As compared to AAS rats, AAR rats showed reduced mRNA levels of the GABA transporter GAT-3 in the CeA but not the BLA. This difference could not be attributed to differential exposure to alcohol since it was similar between AAR and AAS; therefore, it either predates the exposure to alcohol or emerges as vulnerability-specific alcohol-induced adaptation and can therefore be seen as a biological marker of the vulnerability to develop AUD.

These findings are consistent with the previous demonstration of altered expression of several genes involved in GABAergic transmission, including a lower GAT-3 mRNA level, in the amygdala of rats characterised as choosing alcohol over an alternative reward, as well as resisting punishment and quinine adulteration in an instrumental setting^17^. Together these observations suggest that alterations in GABAergic mechanisms in the CeA underlie the compulsive nature of both consummatory and preparatory responses for alcohol, which are otherwise psychologically and neurobiologically dissociable^45, 46^. The suggestion that an altered GABAergic system in the CeA is a biological marker of the vulnerability to AUD is further supported by the presence of similar alterations in humans with AUD^17^.

GABA transporters are involved in the clearance of GABA from the synaptic cleft and therefore play an important role in regulating the extracellular concentration of GABA^20, 39^. Indeed, down-regulation of GAT-3 in rats showing addiction-like behaviours results in an increased GABAergic tone in the CeA, presumably as the result of decreased GABA clearance by these GABA transporters in the synaptic cleft^17^.

Pharmacological activation of GABA_B_ receptors by the agonist baclofen decreases the extracellular release of GABA^23, 24^, and results in a dose-dependent decrease in alcohol intake at the population level, as shown in this and in previous studies^6^. Furthermore, as shown in the present study, baclofen selectively decreased the compulsive drinking of alcohol by AAR rats when challenged by adulteration with quinine. This effect was not attributable to a non-specific effect of baclofen on the aversive properties of quinine because the degree of suppression of drinking following quinine adulteration was not affected by this treatment in AAS rats. This effect was also independent of the influence of baclofen on alcohol drinking by AAR rats at baseline in a two-bottle choice session, as there was no correlation between the suppressant effect on drinking between the two conditions. Together these data suggest that voluntary alcohol drinking and persistent alcohol drinking in the face of negative consequences depend on distinct GABAergic mechanisms. This observation is in agreement with the finding that rats with three addiction-like behaviours^41^ in a multisymptomatic preclinical model^19^ of AUD were specifically sensitive to the motivation- and reinstatement-suppressing effects of baclofen^42^.

Taken together, these results suggest that the decrease in compulsive seeking^42^ and drinking (present study) behaviours shown by individuals with AUD when they are treated with baclofen may be mediated by a normalisation of impaired GABA clearance resulting from the decreased GAT-3 levels. It is important to note, however, that baclofen was administered systemically, thereby precluding any conclusion with regards to its neural locus of action. Further investigations will be necessary to establish whether restoration of GABA levels, specifically and exclusively within the CeA, are necessary and sufficient to decrease compulsive alcohol seeking or drinking.

The psychological consequences of altered GABAergic physiology in the CeA are not fully understood. However, a down-regulation of GAT-3 mRNA levels in the CeA of rats has been associated with increased anxiety^17^, in line with the long established role of the CeA and its GABAergic system in the expression of anxiety related disorders^47, 48^. Considering that anxiety and associated alcohol drinking as a self-medication strategy are important factors of vulnerability in the development of AUD^38,41,49,50^, an altered CeA GABAergic system may represent an endophenotype of this individual vulnerability.

The results of the present study provide new evidence that compulsive alcohol drinking is associated with decreased GAT-3 mRNA levels and is selectively suppressed by treatment with baclofen.

## 5 Acknowledgements

This work was supported by a Programme Grant from the Medical Research Council (MR/NO2530X/1) to BJE and DB and by a Leverhulme Trust Early Career Fellowship to LMP (ECF-2018-713).

## 6 Authors contribution

LMP and DB designed the experiments. LMP, MF, MP and PJC carried-out the behavioural experiment. LMP and ABR performed the molecular procedures. LMP and MF performed the data analysis. LMP and DB wrote the manuscript. BJE gave intellectual input and contributed to the redaction of the manuscript.

## References

1. Geneva: World Heatlh Organization LCB-N-SI. Global status report on alcohol and health 2018. 2018.

2. Anthony JC, Warner LA, Kessler RC. Comparative epidemiology of dependence on tobacco, alcohol, controlled substances, and inhalants: Basic findings from the National Comorbidity Survey. Experimental and Clinical Psychopharmacology. 1994;2(3):244–268.

3. Association AP. The Diagnostic and Statistical Manual of Mental Disorders: DSM 5. bookpointUS; 2013.

4. Everitt BJ, Robbins TW. Drug Addiction: Updating Actions to Habits to Compulsions Ten Years On. Annu Rev Psychol. 2016;67:23–50.

5. Koob GF, Volkow ND. Neurocircuitry of addiction. Neuropsychopharmacology. 2010;35(1):217–238.

6. Colombo G, Gessa GL. Suppressing Effect of Baclofen on Multiple Alcohol-Related Behaviors in Laboratory Animals. Front Psychiatry. 2018;9:475.

7. Koob GF. A role for GABA mechanisms in the motivational effects of alcohol. Biochem Pharmacol. 2004;68(8):1515–1525.

8. Maccioni P, Colombo G. Potential of GABA_B_ Receptor Positive Allosteric Modulators in the Treatment of Alcohol Use Disorder. CNS Drugs. 2019;33(2):107–123.

9. Colombo G, Vacca G, Serra S, Brunetti G, Carai MA, Gessa GL. Baclofen suppresses motivation to consume alcohol in rats. Psychopharmacology (Berl). 2003;167(3):221–224.

10. Lorrai I, Maccioni P, Gessa GL, Colombo G. R(+)-Baclofen, but Not S(-)-Baclofen, Alters Alcohol Self-Administration in Alcohol-Preferring Rats. Front Psychiatry. 2016;7:68.

11. Enoch MA, Hodgkinson CA, Shen PH, et al. GABBR1 and SLC6A1, Two Genes Involved in Modulation of GABA Synaptic Transmission, Influence Risk for Alcoholism: Results from Three Ethnically Diverse Populations. Alcohol Clin Exp Res. 2016;40(1):93–101.

12. Enoch MA, Zhou Z, Kimura M, Mash DC, Yuan Q, Goldman D. GABAergic gene expression in postmortem hippocampus from alcoholics and cocaine addicts; corresponding findings in alcohol-naive P and NP rats. PLoS One. 2012;7(1):e29369.

13. Roberto M, Kirson D, Khom S. The Role of the Central Amygdala in Alcohol Dependence. Cold Spring Harb Perspect Med. 2020.

14. Roberto M, Gilpin NW, Siggins GR. The central amygdala and alcohol: role of gamma-aminobutyric acid, glutamate, and neuropeptides. Cold Spring Harb Perspect Med. 2012;2(12):a012195.

15. Hyytia P, Koob GF. GABA_A_ receptor antagonism in the extended amygdala decreases ethanol self-administration in rats. Eur J Pharmacol. 1995;283(1-3):151–159.

16. Roberts AJ, Cole M, Koob GF. Intra-amygdala muscimol decreases operant ethanol self-administration in dependent rats. Alcohol Clin Exp Res. 1996;20(7):1289–1298.

17. Augier E, Barbier E, Dulman RS, et al. A molecular mechanism for choosing alcohol over an alternative reward. Science. 2018;360(6395):1321–1326.

18. Deroche-Gamonet V, Belin D, Piazza PV. Evidence for addiction-like behavior in the rat. Science. 2004;305(5686):1014–1017.

19. Belin D, Mar AC, Dalley JW, Robbins TW, Everitt BJ. High impulsivity predicts the switch to compulsive cocaine-taking. Science. 2008;320(5881):1352–1355.

20. Jin XT, Galvan A, Wichmann T, Smith Y. Localization and Function of GABA Transporters GAT-1 and GAT-3 in the Basal Ganglia. Front Syst Neurosci. 2011;5:63.

21. Zhou Y, Danbolt NC. GABA and Glutamate Transporters in Brain. Front Endocrinol (Lausanne). 2013;4:165.

22. Spanagel R. Aberrant choice behavior in alcoholism. Science. 2018;360(6395):1298–1299.

23. Delaney AJ, Crane JW, Holmes NM, Fam J, Westbrook RF. Baclofen acts in the central amygdala to reduce synaptic transmission and impair context fear conditioning. Sci Rep. 2018;8(1):9908.

24. Harrison NL. On the presynaptic action of baclofen at inhibitory synapses between cultured rat hippocampal neurones. J Physiol. 1990;422:433–446.

25. Hopf FW, Lesscher HM. Rodent models for compulsive alcohol intake. Alcohol. 2014;48(3):253–264.

26. Vengeliene V, Celerier E, Chaskiel L, Penzo F, Spanagel R. Compulsive alcohol drinking in rodents. Addict Biol. 2009;14(4):384–396.

27. Ahmed SH. Validation crisis in animal models of drug addiction: beyond non-disordered drug use toward drug addiction. Neurosci Biobehav Rev. 2010;35(2):172–184.

28. Carnicella S, Ron D, Barak S. Intermittent ethanol access schedule in rats as a preclinical model of alcohol abuse. Alcohol. 2014;48(3):243–252.

29. Wise RA. Voluntary ethanol intake in rats following exposure to ethanol on various schedules. Psychopharmacologia. 1973;29(3):203–210.

30. Wolffgramm J, Heyne A. From controlled drug intake to loss of control: the irreversible development of drug addiction in the rat. Behav Brain Res. 1995;70(1):77–94.

31. Wolffgramm J. An ethopharmacological approach to the development of drug addiction. Neurosci Biobehav Rev. 1991;15(4):515–519.

32. Spanagel R, Holter SM, Allingham K, Landgraf R, Zieglgansberger W. Acamprosate and alcohol: I. Effects on alcohol intake following alcohol deprivation in the rat. Eur J Pharmacol. 1996;305(1-3):39–44.

33. Giuliano C, Belin D, Everitt BJ. Compulsive Alcohol Seeking Results from a Failure to Disengage Dorsolateral Striatal Control over Behavior. J Neurosci. 2019;39(9):1744–1754.

34. Giuliano C, Pena-Oliver Y, Goodlett CR, et al. Evidence for a Long-Lasting Compulsive Alcohol Seeking Phenotype in Rats. Neuropsychopharmacology. 2018;43(4):728–738.

35. Murray JE, Belin-Rauscent A, Simon M, et al. Basolateral and central amygdala differentially recruit and maintain dorsolateral striatum-dependent cocaine-seeking habits. Nat Commun. 2015;6:10088.

36. Hopf FW, Chang SJ, Sparta DR, Bowers MS, Bonci A. Motivation for alcohol becomes resistant to quinine adulteration after 3 to 4 months of intermittent alcohol self-administration. Alcohol Clin Exp Res. 2010;34(9):1565–1573.

37. Smith JC, Sclafani A. Saccharin as a sugar surrogate revisited. Appetite. 2002;38(2):155–160.

38. Koob GF. The dark side of emotion: the addiction perspective. Eur J Pharmacol. 2015;753:73–87.

39. Guastella J, Nelson N, Nelson H, et al. Cloning and expression of a rat brain GABA transporter. Science. 1990;249(4974):1303–1306.

40. Siciliano CA, Noamany H, Chang CJ, et al. A cortical-brainstem circuit predicts and governs compulsive alcohol drinking. Science. 2019;366(6468):1008–1012.

41. Jadhav KS, Magistretti PJ, Halfon O, Augsburger M, Boutrel B. A preclinical model for identifying rats at risk of alcohol use disorder. Sci Rep. 2017;7(1):9454.

42. Jadhav KS, Peterson VL, Halfon O, et al. Gut microbiome correlates with altered striatal dopamine receptor expression in a model of compulsive alcohol seeking. Neuropharmacology. 2018;141:249–259.

43. Pelloux Y, Everitt BJ, Dickinson A. Compulsive drug seeking by rats under punishment: effects of drug taking history. Psychopharmacology (Berl). 2007;194(1):127–137.

44. Ducret E, Puaud M, Lacoste J, et al. N-Acetylcysteine Facilitates Self-Imposed Abstinence After Escalation of Cocaine Intake. Biol Psychiatry. 2015.

45. Ikemoto S, Panksepp J. Dissociations between appetitive and consummatory responses by pharmacological manipulations of reward-relevant brain regions. Behav Neurosci. 1996;110(2):331–345.

46. Luscher C, Robbins TW, Everitt BJ. The transition to compulsion in addiction. Nat Rev Neurosci. 2020;21(5):247–263.

47. Tye KM, Prakash R, Kim SY, et al. Amygdala circuitry mediating reversible and bidirectional control of anxiety. Nature. 2011;471(7338):358–362.

48. Gilpin NW, Herman MA, Roberto M. The central amygdala as an integrative hub for anxiety and alcohol use disorders. Biol Psychiatry. 2015;77(10):859–869.

49. Koob GF. Brain stress systems in the amygdala and addiction. Brain Res. 2009;1293:61–75.

50. Khantzian EJ. Addiction as a self-regulation disorder and the role of self-medication. Addiction. 2013;108(4):668–669.

